# Painless Na_v_1.7 mutations reveal function-critical residues in the outer vestibule and N-terminus

**DOI:** 10.1101/2025.05.12.653536

**Authors:** Nivedita Sarveswaran, Ichrak Drissi, Samiha Shaikh, Mike Nahorski, Fiona Cusdin, John Linley, C Geoffrey Woods

## Abstract

Chronic pain is a common condition, placing a large cost on society and a heavy burden on the individual. Na_v_1.7 has emerged as a non-redundant and axiomatically important part of human pain pathways. Biallelic loss-of-function (LOF) mutations in *SCN9A*, encoding voltage-gated sodium channel Na_v_1.7, cause the Mendelian disorder Congenital Insensitivity to Pain (CIP). Studying novel missense mutations in this channel has the potential to uncover unrealised functions of critical residues that could enable new strategies for analgesia.

Here we describe detailed functional studies of six *SCN9A* missense variants found in individuals with typical *SCN9A*-CIP: c.224T>C/p.Leu75Pro, c.1025C>T/p.Thr342Met, c.2686C>T/p.Arg896Trp, c.2732G>A/p.Arg911His, c.5059G>C/p.Ala1687Pro, and c.5173G>C/p.Gly1725Arg. While all variants were extremely rare and predicted to alter function, current tools cannot specify the extent or direction of change in function, nor provide information about the mechanism(s) of pathogenicity.

Stable cell lines were generated for each variant, using the isogenic Jump-In™ T-REx™ HEK293 system. We investigated how each variant caused molecular dysfunction using multiple approaches: sequence conservation analysis and 3D modelling to examine the structural importance of the affected residues; automated whole-cell patch clamp electrophysiology to assess channel biophysics; and finally, immunocytochemistry along with cell-surface biotinylation to detect any alterations in subcellular localisation.

Our studies revealed that all mutations reached the cell membrane but caused a complete LOF through small localised structural changes in the extracellular vestibule, ion selectivity filter and N-terminus that abolished sodium conductance. Functional analysis remains the most reliable method of classifying *SCN9A* missense variants and can help to pinpoint important subdomains.

**Online Summary:** Voltage-gated sodium channel Na_v_1.7 is a key modulator of nociceptor excitability and associated with congenital painlessness. Functional analysis and *in silico* modelling reveal critical roles for residues in the extracellular vestibule, ion selectivity filter and unexpectedly, the N-terminus.

## Introduction

Voltage-gated sodium channels (Na_v_ channels) are essential for electrical signalling in neurons. The Na_v_ family comprises nine subtypes, Na_v_1.1 to Na_v_1.9, each with enriched expression in different tissues and sometimes specific subcellular compartments. As a result, pathogenic mutations are associated with disorders of excitability –for example, cardiac arrhythmia or epilepsy caused by Na_v_1.5 or Na_v_1.1 dysfunction respectively (Catterall, 2012; Huang et al., 2017). Na_v_1.7 is enriched in nociceptors, peripheral neurons that detect noxious stimuli and transmit these signals to pain pathways in the central nervous system (Toledo-Aral et al., 1997; Black et al., 2012; Shiers et al., 2021). Mutations in this channel are extremely rare but can cause aberrant pain by gain-of-function (GOF) increasing the open-state probability of the channel, or cause painlessness by loss-of-function (LOF) ablating sodium conductance (Dib-Hajj and Waxman, 2019). Due to the strong human evidence that Na_v_1.7 can act as a bi-directional dial on nociceptor excitability, it has garnered much attention as a target for analgesic drug development (Wulff et al., 2019; Alsaloum et al., 2020).

Eukaryotic Na_v_ channels are produced from a single chain of 1700-2000 amino acids that fold into four structurally homologous domains (DI to DIV), each domain consisting of six α-helical transmembrane segments (S1 to S6) connected by alternating intracellular and extracellular loops. The S5 and S6 segments from each domain form the walls of a central pore which is closed off from the cytoplasm by an intracellular gate – as the S5-S6 segments undergo conformational changes upon channel activation, the intracellular gate is opened. The extracellular S5-S6 loops form a funnel-like vestibule that attracts cations towards the ‘DEKA’ ion selectivity filter (domains: I = D, aspartic acid; II = E, glutamic acid; III = K, lysine; IV = A, alanine), through which only sodium ion flow into the cell is permitted. The S1 to S4 segments of each domain form the outer voltage-sensing modules, with gating charges in the S4 segments shifting vertically through the membrane in response to local changes of membrane potential. Importantly, the intracellular S4-S5 linkers play a critical role in coordinating the movements of these functional modules so that voltage sensor activation is coupled to the opening of the pore (Catterall et al., 2017, 2020).

Of the *SCN9A*-CIP patient mutations that have been functionally assessed *in vitro*, electrophysiological analysis showed that Na_v_1.7 mutants with truncations before the C-terminal tail are completely non-conducting (Cox et al., 2006; Ahmad et al., 2007; He et al., 2018; McDermott et al., 2019). One nonsense mutation, p.Leu1831X, retained partial function as all four voltage sensors, the pore domain and gating components of the α-subunit were intact; only the last 146 residues of the C-terminus were excluded (Emery et al., 2015). Of the nine published missense mutations, four caused a 50-90% reduction in peak current density (Emery et al., 2015; Sun et al., 2020), three blocked sodium conductance completely (Cox et al., 2010; McDermott et al., 2019), and two have not been functionally investigated (Staud et al., 2011; Shorer et al., 2014). In addition to electrophysiology, some groups have used semi-quantitative methods to demonstrate a reduction in total protein expression and plasma membrane localisation caused by *SCN9A* mutations (Cox et al., 2010; Sun et al., 2020).

Functional evidence showing a suspected variant causes molecular dysfunction is vital for clinicians to make an accurate genetic diagnosis (Duzkale et al., 2013; MacArthur et al., 2014). This is particularly important in ultra-rare, autosomal recessive disorders such as *SCN9A*-CIP so that related carriers can receive appropriate genetic counselling (Schon et al., 1993). In this study, we identified six missense mutations in *SCN9A*-CIP patients. Their unique clinical phenotype, and the presence of these missense mutations in trans with a second *SCN9A* mutation, provided strong grounds for pathogenicity. We employed multiple molecular assays to investigate the effect of these variants on channel biophysics and subcellular distribution, including cell membrane localisation. In addition, we examined Na_v_1.7 atomic structures to understand how these localised changes could impact channel function.

## Methods

### Clinical and Genetic Testing

The six individuals in this study were ascertained in a supra-regional Paediatric Neurology/Clinical Genetics clinic for the diagnosis and management of Mendelian disorders of painlessness (Schon et al., 1993). Initial clinical diagnoses of *SCN9A*-CIP were made in all six. Genetic testing of the affected child and unaffected parents was instigated after discussion with the parents and their consent. Genetic testing was carried out by NHS exome or genome of the proband, then filtered for known genes associated with painlessness. Families also consented to research studies; research ethics was granted by the Cambridge East Research Ethics Committee (12/EE/40486). All variants were confirmed by Sanger sequencing and segregation was assessed by Sanger sequencing of the parents. The pathogenicity of patient variants in Na_v_1.7 was evaluated based on evidence from previously reported mutations and predictions using the PolyPhen-2 tool configured to the HumVar-trained model (Adzhubei et al., 2013). Allele frequency data was collected from the gnomAD database (Chen et al., 2024).

### Bioinformatics

For conservation analysis, consensus amino acid sequences for human Na_v_1.1 to Na_v_1.9 and Na_v_1.7 homologues across species were obtained from UniProt (UniProt Consortium, 2021). Multiple sequence alignments were generated using the EMBL-EBI Clustal Omega tool (Sievers et al., 2011) and illustrated using Jalview (Waterhouse et al., 2009). For structural modelling, we used the PDB model 7W9K from the cryo-EM resolved structure of human Na_v_1.7 (Huang et al., 2022), loaded into Chimera software (Pettersen et al., 2004) to visualise the location of patient mutations. This structure represents the 5N11L isoform of Na_v_1.7 – the neonatal vs adult splice variants at exon 5 (5N vs 5A) encode alternative amino acids at positions 201 and 206 within the DI:S3-S4 intracellular linker. The longer exon 11 variant (11L) adds a short peptide sequence within the domain I-II intracellular loop compared to the shorter (11S) isoforms (Raymond et al., 2004). None of our mutations affect these residues, however, the amino acid changes we report are based on numbering from the 11S isoform; these discrepancies are noted where relevant.

### Plasmid Cloning

The wild-type *SCN9A* plasmid was kindly gifted by Fiona Cusdin (MedImmune, UK). The insert encodes the 5N11S isoform of Na_v_1.7 and was codon-optimised for mammalian expression. It includes a FLAG epitope tag and PreScission protease recognition sequence at the N-terminus of *SCN9A*, followed by a short flexible linker, thrombin cleavage site and poly-histidine tag at the C-terminus. The insert was cloned into the pJTI™ R4 Dest CMV-TO pA vector, which is necessary for generating stable cell lines using the Jump In™ T-REx™ HEK 293 system as detailed in the next section. Patient mutations were introduced into the *SCN9A* plasmid using the Quikchange II XL site-directed mutagenesis kit (Agilent Technologies). Plasmids were grown using MAX Efficiency Stbl2 Competent cells (Invitrogen) according to a recommended protocol (Feldman and Lossin, 2014). Each new batch of plasmids was sequence verified before use in experiments.

### Stable Cell Line Generation

The Jump-In™ T-REx™ HEK293 system (Thermo Scientific) was used to generate isogenic cell lines with stable integration and inducible expression of Na_v_1.7. The parental cell line has been engineered to include three key features (Supplementary Figure 1A). Firstly, it carries a single retargeting acceptor site ‘*attP*’ which undergoes homologous recombination with the donor site ‘*attB*’ in the pJTI™ R4 Dest CMV-TO pA partner plasmid, but only when R4 integrase is co-transfected. Secondly, a promoter-less *NeoR* gene, conferring resistance to selection antibiotic G418, is placed alongside the *attP* site in the parental cell line which is only activated when the EF1α promoter is introduced by recombination with the partner plasmid. Lastly, the parental cell line stably expresses the tetracycline repressor protein which binds to the tetracycline operator sequence downstream of the CMV promoter. This prevents gene transcription until it is displaced by doxycycline added to the cell culture medium.

**Supplementary Figure 1:**
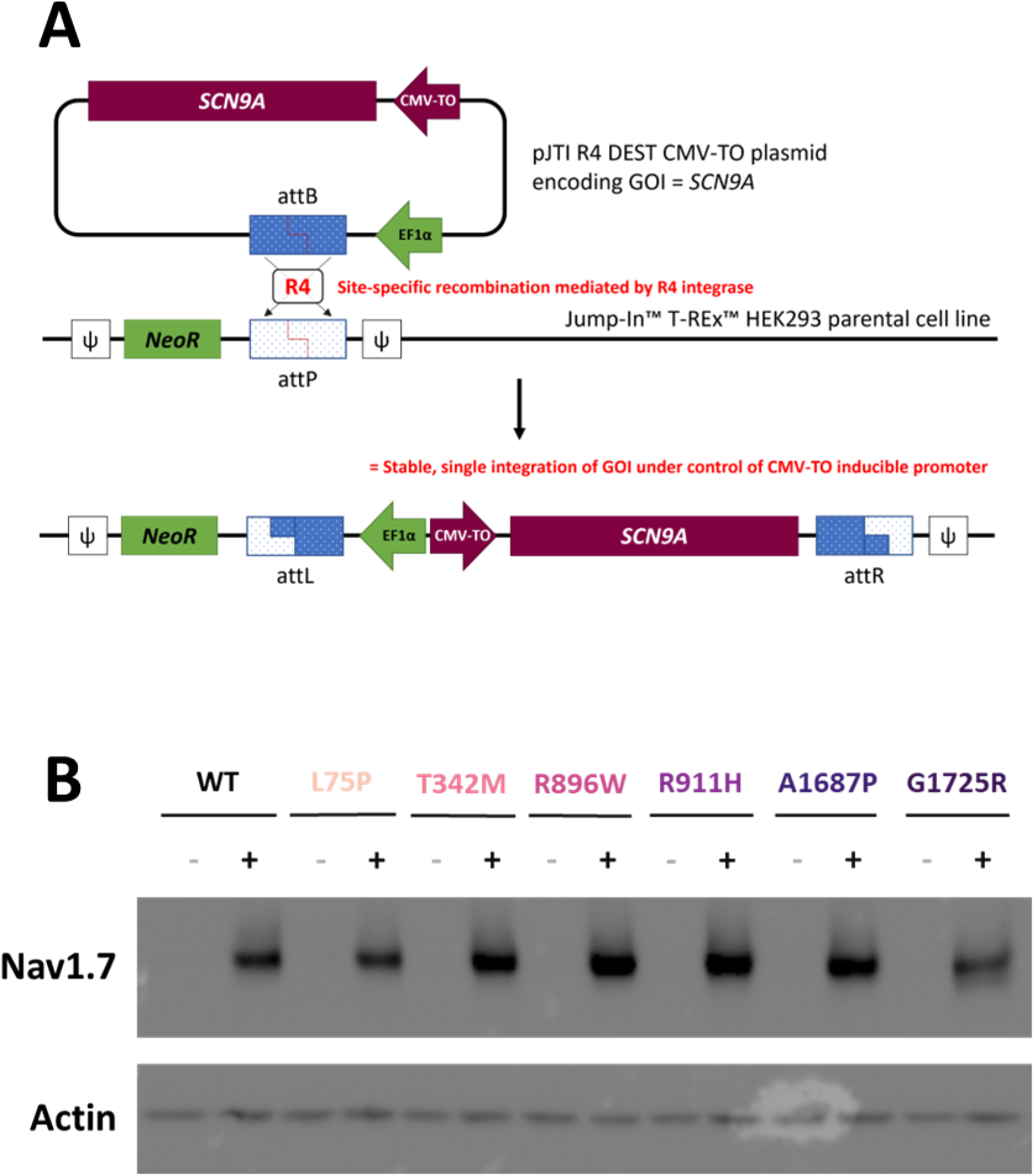
Generation and validation of Na_v_1.7 stable cell lines. (A) Schematic overview of the pJTI™ R4 Dest CMV-TO plasmid encoding the gene of interest (GOI) *SCN9A* driven by a CMV-TO promoter and also containing the EF1α promoter. After undergoing homologous recombination with the Jump-In™ T-REx™ HEK293 parental cell line, assisted by R4 integrase, the EF1α promoter now drives constitutive *NeoR* expression. Thus, only cells with successful integration at this site would be resistant to Geneticin treatment, resulting in polyclonal but isogenic stable cell lines. (B) Western blot demonstrating inducible expression of Na_v_1.7 in our stable cell lines where (−) and (+) indicate doxycycline absent vs doxycycline present in cell media, respectively.

Stable cell lines were generated for wild-type Na_v_1.7 and the six novel CIP patient mutations, as well as a previously characterised CIP mutation p.Ala1236Glu (Emery et al., 2015) and emGFP, to use as controls in electrophysiology experiments. Cell lines were maintained in Dulbecco’s modified Eagle’s medium with 10% dialysed foetal bovine serum, 1% non-essential amino acids, 5 µg/ml Blasticidin S HCl and 1 mg/ml Geneticin (all Gibco). To induce Na_v_1.7 expression, cell culture medium was supplemented with 1 µg/ml doxycycline hydrochloride for 16-24 hours (Supplementary Figure 1B).

### Electrophysiology

Automated electrophysiology was performed on the Sophion QPatch 16 platform (Sophion Bioscience), which can accommodate up to 16 cells for simultaneous patch clamp recording to conduct whole-cell patch clamp electrophysiology on our Na_v_1.7 cell lines. The intracellular solution contained (in mM): 10 NaCl, 140 CsF, 2 EGTA and 10 HEPES, pH 7.3 by CsOH titration and adjusted to 305-315 mOsm. The extracellular solution contained (in mM): 140 NaCl, 2 KCl, 2 CaCl_2_, 1 MgCl_2_ and 10 HEPES, pH 7.4 by NaOH titration and adjusted to 295-305 mOsm. The stable cell lines were detached using Accutase (Gibco), then spun down and gently re-suspended in FreeStyle Expression Medium (Gibco) supplemented with 20 mM HEPES and trypsin inhibitor. The suspension was filtered through a 40 µM cell strainer before being applied to compound plates used with the QPatch system.

Series resistance compensation (70%) was utilised to avoid voltage drop artefacts and seal resistance was >1GΩ. Cells were voltage clamped at −80 mV and a continuous voltage pulse was applied every 5s to monitor the Na_v_1.7 current (100 ms at −120 mV followed by 10 ms at 0 mV and −120 mV for 100 ms; sampled at 10 KHz). Voltage dependence of activation and current/voltage relationship for Na_v_1.7 WT and mutant constructs was measured using 200 ms voltage steps from a holding potential of −120 mV up to +20 mV in 5 mV increments.

Raw data was handled by Sophion Analyzer software before being exported to Microsoft Excel for analysis. Currents generated from the voltage dependence of activation protocol were transformed into conductance values for each cell using the following equation, where *G* represents conductance, *I* represents current, *V* is the membrane potential measured at each voltage step and *V_rev_* is the reversal potential per cell. The reversal potential was calculated on a cell-by-cell basis by fitting a linear regression across the currents produced at voltage steps 0 to 20 mV, then calculating the voltage at which there is no net flow of sodium ions.

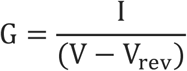

To determine the voltage dependence of activation, conductance at each voltage step was normalised to be expressed as a fraction of the peak amplitude. Boltzmann curves were fit using the equation below, where *G/G_max_* represents the available fraction of normalised conductance, *V_1/2_* is the half-maximal activation potential, *V* is the voltage step and *k* is the slope factor:

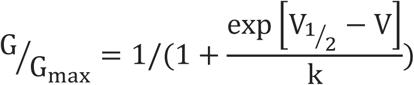

Statistical analysis, curve-fitting and data plotting were done in GraphPad Prism. The peak currents from Na_v_1.7 stable cell lines were compared using the Brown-Forsythe and Welch one-way ANOVA, with Dunnett T3 correction for multiple comparisons. Results in the main text are depicted as the mean average ± standard error of the mean (SEM) across three experimental replicates.

### Immunocytochemistry

Stable cell lines were plated onto poly-L-lysine coated coverslips in the presence of doxycycline to induce Na_v_1.7 expression. Cells were fixed with 4% paraformaldehyde, permeabilised with 0.1% Triton X-100 and blocked using 10% normal donkey serum and 10% bovine serum albumin – all in PBS. Cells were incubated with primary antibodies at 4°C overnight, followed by incubation with secondaries at room temperature for 1 hour. Finally, they were mounted onto slides using Prolong Diamond Antifade Mountant with DAPI (Invitrogen). Antibodies used were: mouse anti-Na_v_1.7 (RRID:AB_10808664, 1:200), rabbit anti-KDEL (RRID:AB_2819147, 1:500), donkey anti-rabbit AF488 (RRID:AB_2535792, 1:1000) and donkey anti-mouse AF546 (RRID:AB_11180613, 1:1000). Images were acquired using the LSM880 confocal microscope (Zeiss) with a 63x magnification, planar-apochromatic, 1.4NA oil immersion lens. Excitation by Diode 405 nm, Argon 488 nm and Diode 561 nm lasers were used to detect DAPI staining, AF488 and AF546 fluorophores respectively. Images were collected across two independent cultures.

### Cell-Surface Biotinylation

In preliminary experiments involving transient transfection, HEK293 cells were plated at 1.5 × 10^5^ cells/ml per 100 mm dish, then transfected the next day using a 3:1 ratio of FuGENE HD (Promega) transfection reagent to *SCN9A* plasmid DNA. In later experiments involving the stable cell lines, *SCN9A* expression was induced the day after plating by adding doxycycline to the media. In both cases, biotinylation was performed after 24 hours of gene expression, using the Pierce Cell Surface Protein Isolation Kit (Thermo Scientific). Briefly, cells were washed twice with ice-cold PBS and then incubated with 0.25 mg/ml EZ-Link Sulfo-NHS-SS-Biotin per dish, for 30 minutes at 4°C. The reaction was stopped with the provided quenching solution containing 100 mM glycine. Cells were then pelleted and washed twice in ice-cold TBS, re-suspended in 500 μl lysis buffer, and disrupted by sonication. Cell lysis proceeded on ice for 30 minutes with periodic vortexing. A 100 μl aliquot of lysed cells was set aside as the “total” fraction. The remainder of the lysed cells were added to a NeutrAvidin agarose column and incubated for 1 hour at room temperature to bind the biotinylated surface proteins. After 3 washes with wash buffer, the column was heated for 5 minutes at 95°C and biotinylated proteins (“surface” fraction) were eluted in 400 μl NuPAGE LDS sample buffer (Invitrogen) containing 50mM dithiothreitol. EDTA-free Protease Inhibitor Mini Tablets (Thermo Scientific) were added to the supplied lysis and wash buffers. Three experimental replicates were performed in the transient system for all variants except p.Ala1687Pro due to this variant being identified later in the project, however, we did study this mutation in triplicate in the stable cell line system. Due to limited resources during the COVID-19 pandemic, we could only perform two experimental replicates on the remaining stable cell lines.

### Western Blot

Input samples from the biotinylation experiments were prepared in NuPAGE LDS sample buffer with 2.5% β-mercaptoethanol and incubated for 10 minutes at 70°C; these conditions were optimised to produce the tightest bands. Proteins were separated using NuPAGE 3-8% Tris-Acetate gels and transferred onto PVDF membranes at 20V for 13 minutes using the iBlot2 dry transfer system. Membranes were blocked with 5% milk, 1% bovine serum albumin, 0.1% tween-20 buffer in PBS. Antibodies used were: mouse anti-Na_v_1.7 (RRID:AB_10808664, 1:500), rabbit anti-Na^+^K^+^ATPase (RRID:AB_1310695, 1:10000), mouse anti-β-actin (RRID:AB_476697, 1:5000), rabbit anti-calnexin (RRID:AB_2040667, 1:2500), HRP-linked anti-mouse IgG (RRID:AB_330924, 1:1000), and HRP-linked anti-rabbit IgG (RRID:AB_2099233, 1:1000). Bands were visualised using the Amersham ECL Prime Western Blotting Detection Reagent (GE Healthcare) and ChemiDoc MP Imaging System and Image Lab software (Bio-Rad).

## Results

### Novel *SCN9A* missense variants identified in CIP patients

Six patients received a provisional diagnosis of CIP due to their impaired pain sensation with evidence of repeated painless injuries. Given their normal temperature regulation, cognitive development, intestinal function, and absence of *Staphylococcus aureus* infections but the presence of anosmia, the clinical presentation pointed to LOF *SCN9A* mutations as the cause. Genetic testing confirmed this was the case as known or predicted pathogenic variants were found in both *SCN9A* alleles (see Table 1), consistent with an autosomal recessive mode of inheritance. Six novel missense variants were identified (amino acid annotation based on the 11S isoform of Na_v_1.7): p.Leu75Pro, p.Thr342Met, p.Arg896Trp, p.Arg911His, p.Ala1687Pro, p.Gly1725Arg.

**Table 1:**
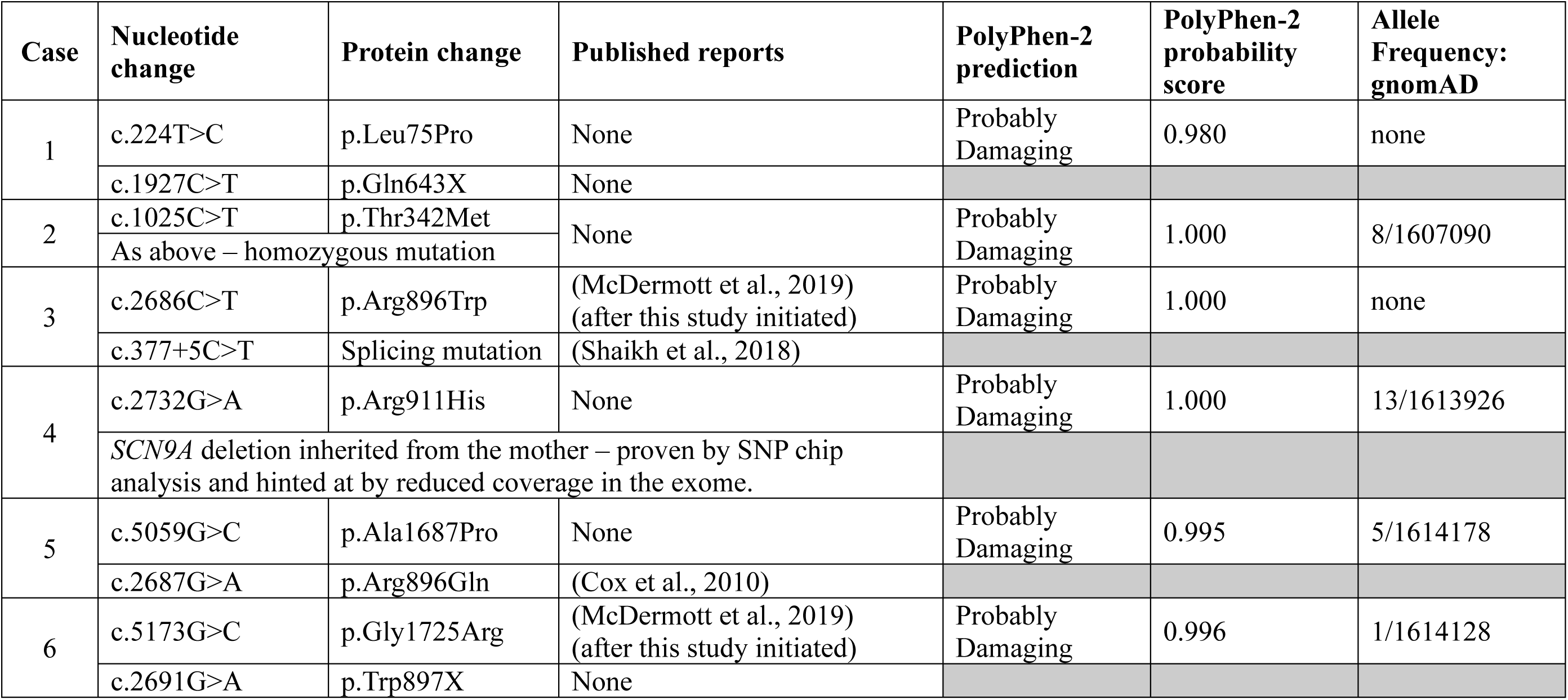
*SCN9A* variants detected in CIP patients.

| Case | Nucleotide change | Protein change | Published reports | PolyPhen-2 prediction | PolyPhen-2 probability score | Allele Frequency: gnomAD |
| --- | --- | --- | --- | --- | --- | --- |
| 1 | c.224T>C | p.Leu75Pro | None | Probably Damaging | 0.980 | none |
|  | c.1927C>T | p.Gln643X | None |  |  |  |
| 2 | c.1025C>T | p.Thr342Met | None | Probably Damaging | 1.000 | 8/1607090 |
|  | As above – homozygous mutation |  |  |  |  |  |
| 3 | c.2686C>T | p.Arg896Trp | (McDermott et al., 2019)<br>(after this study initiated) | Probably Damaging | 1.000 | none |
|  | c.377+5C>T | Splicing mutation | (Shaikh et al., 2018) |  |  |  |
| 4 | c.2732G>A | p.Arg911His | None | Probably Damaging | 1.000 | 13/1613926 |
|  | SCN9A deletion inherited from the mother – proven by SNP chip analysis and hinted at by reduced coverage in the exome. |  |  |  |  |  |
| 5 | c.5059G>C | p.Ala1687Pro | None | Probably Damaging | 0.995 | 5/1614178 |
|  | c.2687G>A | p.Arg896Gln | (Cox et al., 2010) |  |  |  |
| 6 | c.5173G>C | p.Gly1725Arg | (McDermott et al., 2019)<br>(after this study initiated) | Probably Damaging | 0.996 | 1/1614128 |
|  | c.2691G>A | p.Trp897X | None |  |  |  |

All six variants were predicted to be “Probably Damaging” by PolyPhen-2 – this tool is trained to distinguish rare deleterious mutations from common polymorphisms that may cause mild changes in function by evaluating sequence conservation across protein families and scoring the biochemical impact of amino acid substitutions on local structural features (Adzhubei et al., 2013). All variants were present at extremely low allele frequencies in gnomAD, a global genome aggregation database, in the range expected for a very rare recessive Mendelian disorder. At the time of sequencing these variants were also previously unreported as pathogenic, but the p.Arg896Trp and p.Gly1725Arg mutations have since been functionally assessed by another group (McDermott et al., 2019).

### *In silico* analysis illustrates the structural importance of the affected Na_v_1.7 residues

Sequence analysis of Na_v_1.7 across species, from fruit flies to large mammals, shows complete conservation of the native residues that were mutated in our CIP patients (Figure 1A). For four of the affected residues – Leu75, Thr342, Arg 911 and Ala1687 – there is also complete conservation within the human Na_v_ family. The remaining two affected residues – Arg896 and Gly1725 – are fully conserved from Na_v_1.1 to Na_v_1.8 but are substituted in Na_v_1.9 to His896 and Ala1725. These findings suggest that to maintain normal channel function, there is minimal tolerance for substitutions at the affected locations.

**Figure 1:**
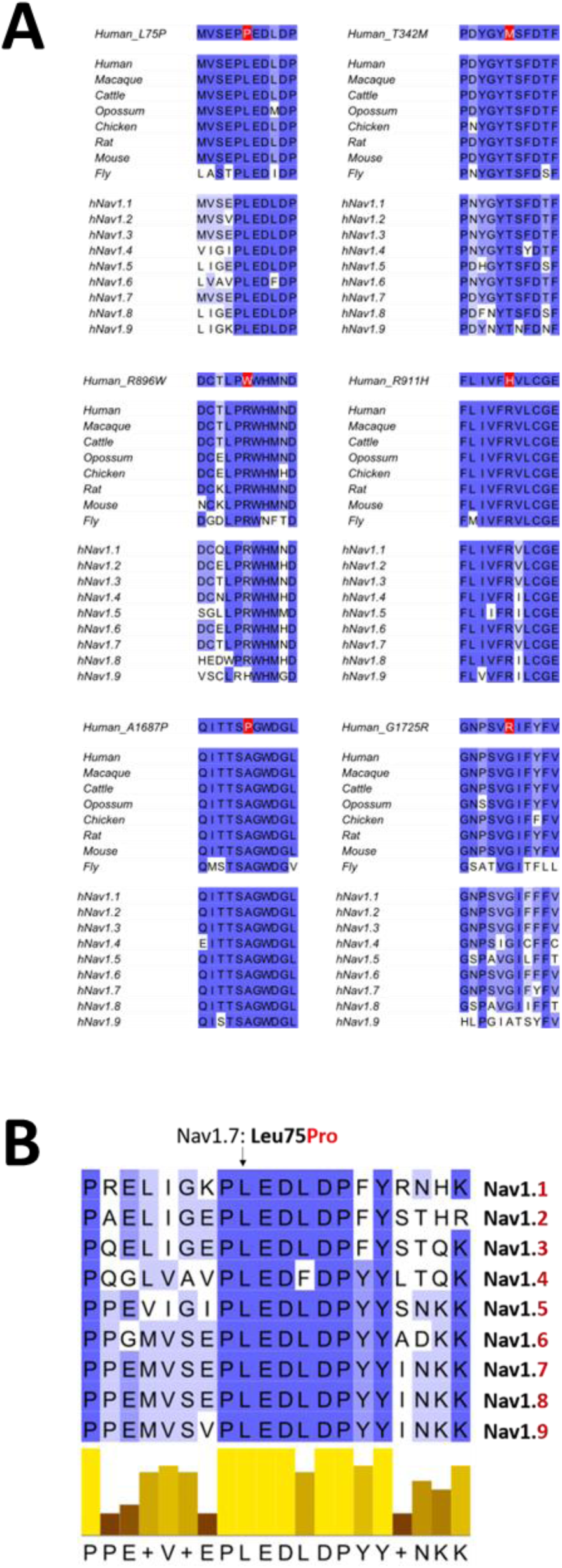
Sequence conservation analysis of native residues across Na_v_1.7 homologues. Multiple sequence alignments generated using the EBI-EMBL Clustal Omega tool to assess tolerance for substitutions of the affected residues in our CIP patients. (A) Na_v_1.7 orthologues across species and across the human Na_v_ channel family. (B) The N-terminal “PLEDLDPYY” motif across the human Na_v_ channel family, where p.Leu75Pro Na_v_1.7 mutation is located.

Based on sequence-structure annotation (Klugbauer et al., 1995) and the recently solved cryo-EM structure of human Na_v_1.7 (Huang et al., 2022), the new Na_v_1.7 mutations were mapped onto their 3D locations and partner residues that form hydrogen bonds via their sidechains were revealed (Figure 2 and Supplementary Video 1). Leu75 is in the intracellular N-terminal tail, within a 9 amino acid motif (PLEDLDPYY) that is highly conserved across Na_v_1.7 orthologues and the human Na_v_ family (Figure 1B), compared to the rest of the N-terminal sequence. Thr342 is in the domain I:S5-S6 pore loop, partially buried under the extensive loops that form the outer vestibule, forming hydrogen bonds with Asp338 – a location only occupied by aspartic acid or asparagine across all Na_v_1.7 homologues.

**Figure 2:**
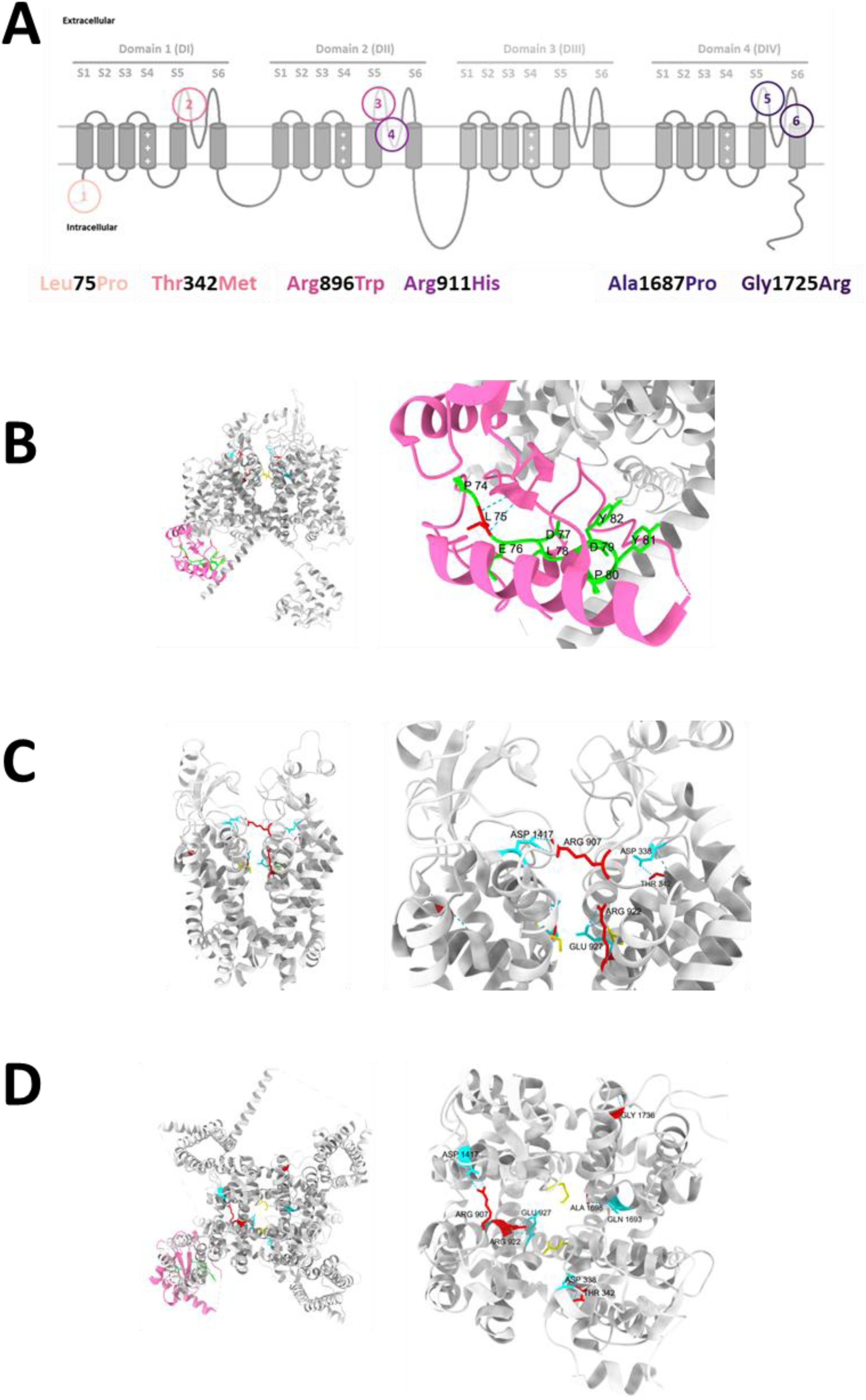
Structural context of native residues affected by *SCN9A*-CIP mutations. (A) Topology of Na_v_1.7 highlighting the six transmembrane segments per domain and the locations of the CIP mutations. (B – D) PDB entry 7W9K for human Na_v_1.7 was rendered in ChimeraX, which represents an exon 11L splice variant. The native residues affected by our CIP mutations are coloured in red and labelled Leu75, Thr342, Arg907 (=Arg896), Arg922 (=Arg911), Ala1698 (=Ala1687) and Gly1736 (=Gly1725). Partner residues Thr338, Glu927, Asp1417 and Gln1693 forming hydrogen bonds via their sidechains are coloured in cyan. The ion selectivity filter consists of Asp361, Glu927, Lys1406 and Ala1698 – residues unaffected by our mutations are coloured in yellow. (B) Side view of the channel with zoom in on the N-terminal tail coloured in pink, the “PLEDLDPYY” motif in green and Leu75 in red. (C) Side view of the pore domain only, with zoom in highlighting Thr342, Arg907 and Arg922. (D) Top view of the channel, with zoom in highlighting all mutations except Leu75.

<u>**Supplementary Video 1: Na_v_1.7 3D structure highlighting intramolecular interactions of native residues affected by *SCN9A*-CIP mutations**</u>

PDB entry 7W9K for human Na_v_1.7 was rendered in ChimeraX, which represents an exon 11L splice variant. The native residues affected by our CIP mutations are coloured in red and labelled Leu75, Thr342, Arg907 (=Arg896), Arg922 (=Arg911), Ala1698 (=Ala1687) and Gly1736 (=Gly1725). Partner residues Thr338, Glu927, Asp1417 and Gln1693 forming hydrogen bonds via their sidechains are coloured in cyan. The ion selectivity filter consists of Asp361, Glu927, Lys1406 and Ala1698 – residues unaffected by our mutations are coloured in yellow. The N-terminal tail (residues 1-124) is coloured in pink, excluding the “PLEDLDPYY” motif in green which is highly conserved across Na_v_1.7 homologues and holds our Leu75 mutation.

Arg896 and Arg911 are in the domain II:S5-S6 pore loop but slightly more exposed to the extracellular space. Arg896 bridges DII-DIII pore loops by forming a salt bridge with Asp1406 (Asp1417 in exon 11L splice variants). This aspartic acid is fully conserved across all Na_v_1.7 homologues. Arg911 is directly jutting into the ion permeation pathway and forms a salt bridge with Glu916 (Glu927 in exon 11L splice variants) below it, the second amino acid of the ion selectivity filter. Ala1687 is within the core of the channel and is the final residue of the sodium ion selectivity filter, forming a hydrogen bond with Gln1682 which is mostly conserved among Na_v_1.7 homologues apart from a substitution to glutamic acid in human Na_v_1.4. Gly1725 is the top-most residue of the alpha-helical domain IV:S6 transmembrane segment.

### Electrophysiological analysis revealed all mutant channels were non-conducting

We conducted biophysical profiling of the mutant channels using our stable cell lines on an automated patch clamp platform. Doxycycline-induced expression of the wild-type and mutant Nav1.7 cell lines was confirmed by western blot (Supplementary Figure 1). For negative controls, we used an additional stable cell line with induced expression of emGFP, and the wild-type Na_v_1.7 cell line without doxycycline induction. We also made a stable cell line with the previously characterised partial LOF mutant, p.Ala1236Glu (Emery et al., 2015) to check if we could capture small currents. When Na_v_1.7 expression was induced, the wild-type cell line displayed a normal I-V curve, with half-maximal activation close to reported values in the literature (V_50_ = −30.5 ± 0.2 mV; k = 3.0 ± 0.2; n = 20 cells) (Figure 3A) (Estacion et al., 2013; Blesneac et al., 2018). In contrast, all other recorded cell lines demonstrate negligible current from which a meaningful V_50_ for voltage-dependence of activation could be determined (Figure 3B).

**Figure 3:**
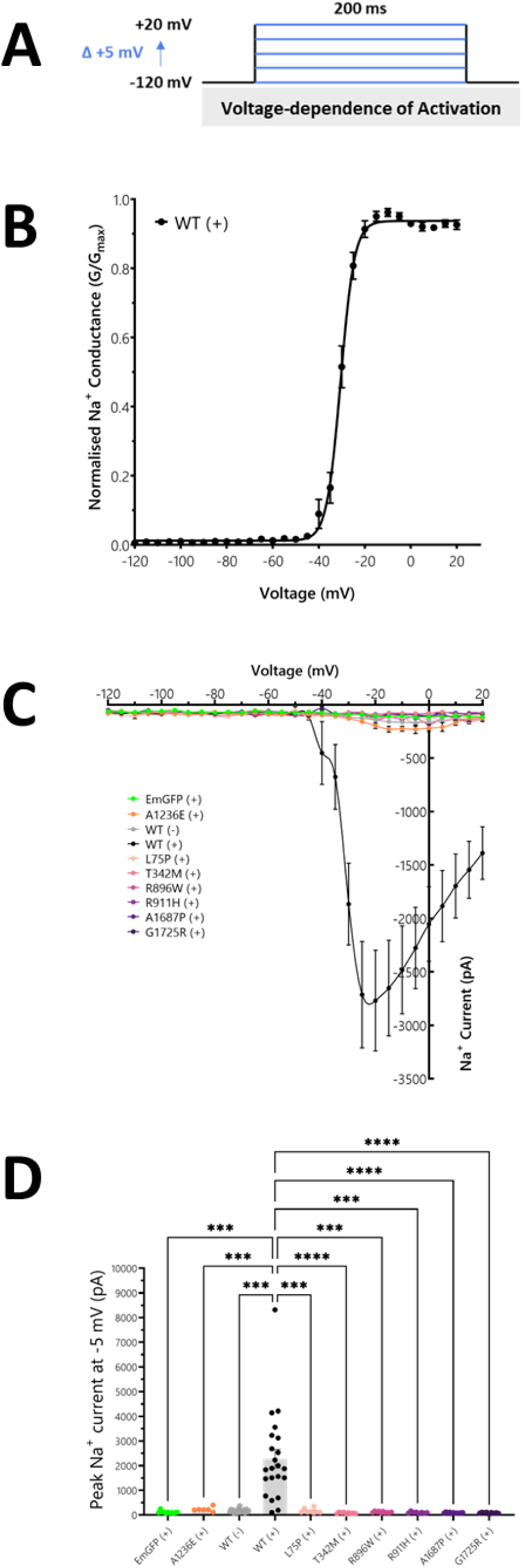
Whole-cell patch clamp analysis of Na_v_1.7 stable cell lines. Na_v_1.7 Jump-In™ T-REx™ HEK293 stable cell lines were evaluated by whole-cell patch clamp on the automated QPatch platform. (+) and (−) symbols indicate whether doxycycline was present or absent in media to induce gene expression. (A) Voltage-dependence of activation protocol comprised of measuring currents during 200 ms voltage steps from a holding potential of −120 mV up to +20 mV in 5 mV increments. (B) Voltage-dependence of activation curve for the wild-type (WT) Na_v_1.7 stable cell line determined V_50_ = −30.5 ± 0.2 mV; k = 3.0 ± 0.2 (n = 20 cells). (C) Current-voltage relationships for all stable cell lines, including negative controls emGFP(+) and WT(−), plus the A1236E(+) Na_v_1.7 mutant as a positive control for partial current. (D) Summary of peak current amplitude at −5 mV across the stable cell lines; these results show the absence of current produced from all new Na_v_1.7 mutations, statistically significant (p <0.001) compared to wild-type Na_v_1.7 as determined by one-way ANOVA with Dunnett T3 correction for multiple comparisons.

For statistical comparison of peak current amplitudes across the different cell lines, we took the average value at −5 mV, as this was where the largest currents were detected for the partial LOF mutant p.Ala1236Glu and our new CIP variants (Figure 3C and Table 2). Taking the negative controls as our baseline for no Nav channel current, the emGFP-induced cells and wild-type Na_v_1.7 non-induced cells exhibited average currents of 117.7 ± 15.0 pA and 168.8 ± 15.0 pA respectively. In contrast, the cells with wild-type Na_v_1.7 induction produced robust sodium currents, averaging 2275.8 ± 380.1 pA at −5 mV. The average current at −5mV for p.Ala1236Glu mutant cells was slightly higher than negative controls, at 231.0 ± 38.8 pA. However, all the new CIP variant cell lines produced negligible sodium current, similar to or even lower than the negative controls – ranging from 167.9 ± 33.3 pA for the p.Leu75Pro cells to 82.1 ± 9.3 for the p.Thr342Met cells. These results confirm that the new *SCN9A* missense mutations from our CIP patients cause complete LOF in Na_v_1.7.

**Table 2:**
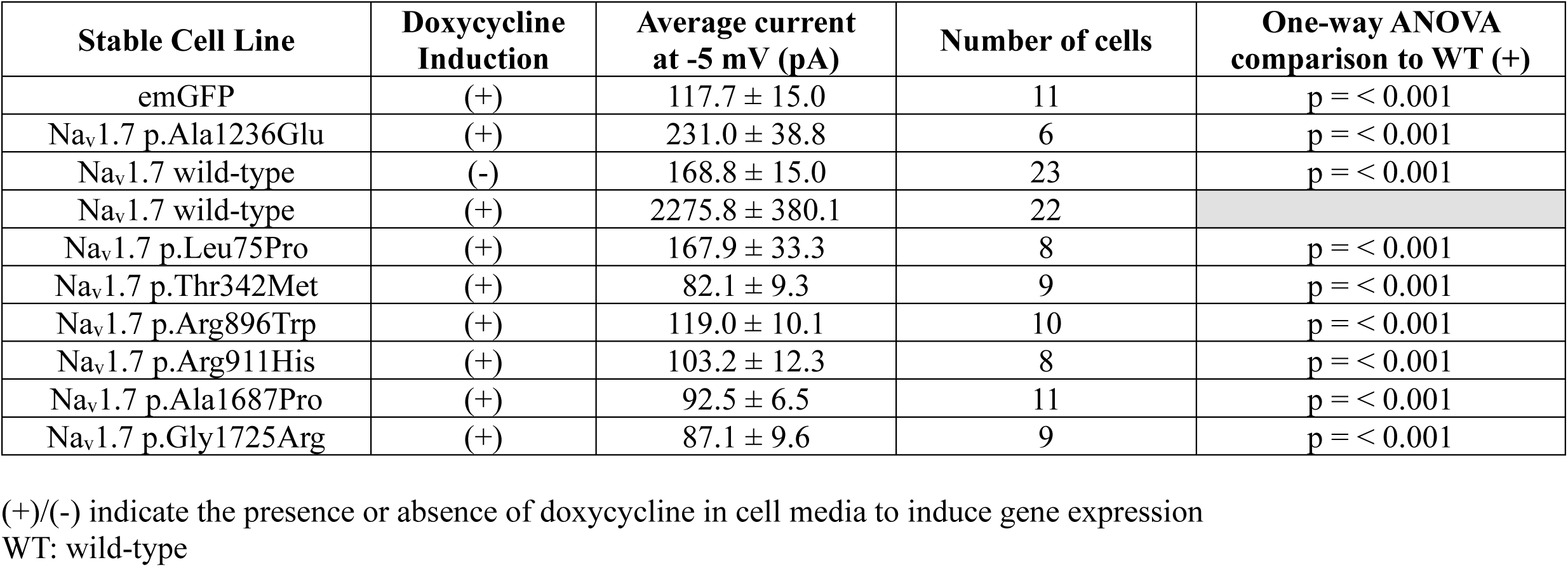
Whole-cell currents produced from Na_v_1.7 stable cell lines.

| Stable Cell Line | Doxycycline Induction | Average current at -5 mV (pA) | Number of cells | One-way ANOVA comparison to WT (+) |
| --- | --- | --- | --- | --- |
| emGFP | (+) | 117.7 ± 15.0 | 11 | p = < 0.001 |
| Nav1.7 p.Ala1236Glu | (+) | 231.0 ± 38.8 | 6 | p = < 0.001 |
| Nav1.7 wild-type | (-) | 168.8 ± 15.0 | 23 | p = < 0.001 |
| Nav1.7 wild-type | (+) | 2275.8 ± 380.1 | 22 |  |
| Nav1.7 p.Leu75Pro | (+) | 167.9 ± 33.3 | 8 | p = < 0.001 |
| Nav1.7 p.Thr342Met | (+) | 82.1 ± 9.3 | 9 | p = < 0.001 |
| Nav1.7 p.Arg896Trp | (+) | 119.0 ± 10.1 | 10 | p = < 0.001 |
| Nav1.7 p.Arg911His | (+) | 103.2 ± 12.3 | 8 | p = < 0.001 |
| Nav1.7 p.Ala1687Pro | (+) | 92.5 ± 6.5 | 11 | p = < 0.001 |
| Nav1.7 p.Gly1725Arg | (+) | 87.1 ± 9.6 | 9 | p = < 0.001 |

(+)/(-) indicate the presence or absence of doxycycline in cell media to induce gene expression
WT: wild-type

### Immunocytochemistry showed wild-type and mutant channels exhibit a similar subcellular distribution

We used confocal microscopy to examine the subcellular localisation of the Na_v_1.7 mutant channels in our stable cell lines, primarily to see whether channel trafficking to the cell membrane could explain the lack of sodium current. Both wild-type and mutant Na_v_1.7 channels exhibited a similar diffuse intracellular distribution in this HEK293 cell background (Figure 4). When combined with an endoplasmic reticulum (ER) marker, there was no obvious overlap in staining that would suggest retention in the ER, as has been observed for other misfolded channels (Manganas et al., 2001; Marangoni et al., 2011). Despite the lack of obvious differences in subcellular distribution, we could not detect (and thus assess) Na_v_1.7 localisation at the cell membrane, even when using sodium-potassium ATPase or wheat germ agglutinin to demarcate this region (data not shown). However, this was not surprising as a Na_v_1.7 “cell membrane ring” has not yet been reported using an immunocytochemistry approach at this resolution in HEK293 cells. Therefore, we looked for an alternative assay to measure Na_v_1.7 localised at the cell membrane.

**Figure 4:**
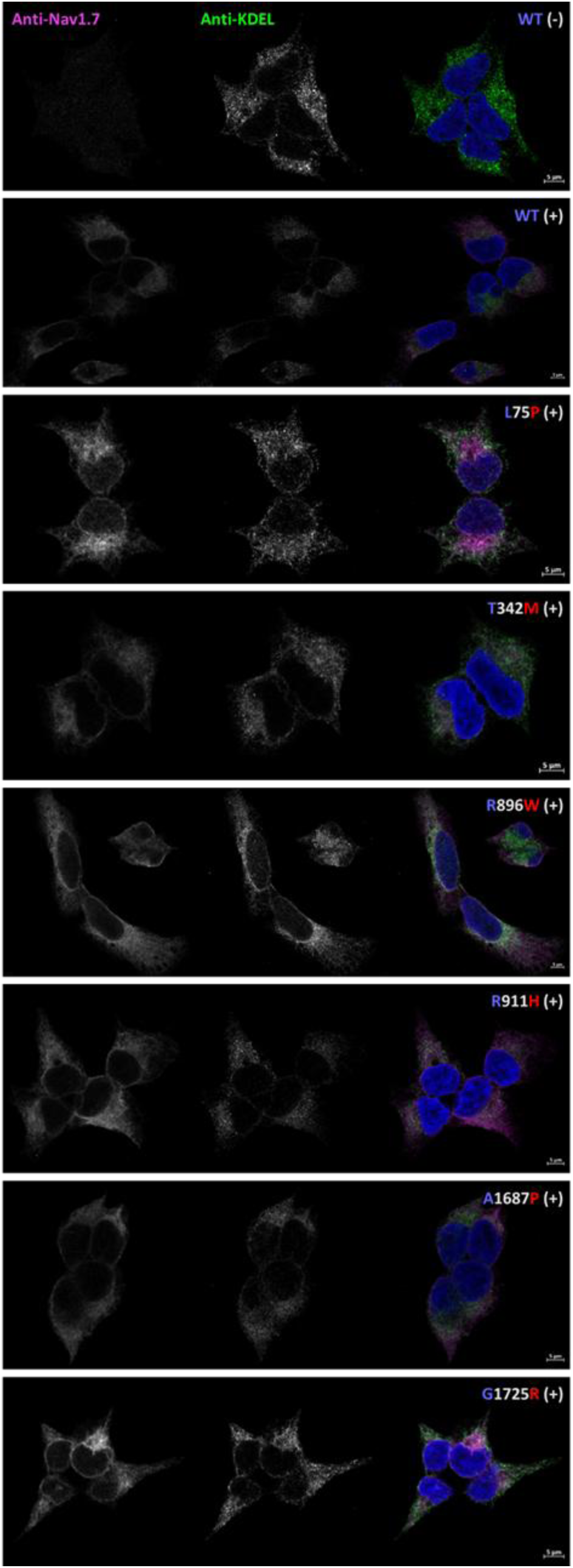
Subcellular distribution of Nav1.7 in stable cell lines. Na_v_1.7 Jump-In™ T-REx™ HEK293 stable cell lines were fixed with paraformaldehyde, permeabilised with Triton X-100, blocked with bovine serum albumin and donkey serum, then stained with anti-Na_v_1.7 and anti-KDEL as an endoplasmic reticulum (ER) marker. Na_v_1.7 and KDEL signals are pseudocoloured in magenta and green respectively. Wild-type and mutant Na_v_1.7 channels exhibited a diffuse intracellular pattern which extended to the tips of the cells but did not display any obvious overlap with the ER.

### Cell-surface biotinylation indicated all mutant channels reached the cell membrane

Cell-surface biotinylation uses the membrane-impermeable reagent sulfo-NHS-SS-biotin to label proteins localised at the cell surface. This assay requires the proteins of interest to have extracellular-facing Lysine residues that react with the N-hydroxysuccinimide group; we expected this to be successful for Na_v_1.7 given previous reports using this approach (Laedermann et al., 2013; Dustrude et al., 2013).

In preliminary experiments using transiently transfected HEK293 cells (n=3), we found 5 mutant channels reached the cell membrane – a representative blot is shown in (Figure 5A). This assay was then repeated on our stable cell lines, by which time the sixth mutation, p.Ala1687Pro, had been identified in a new *SCN9A*-CIP patient. Due to the limited availability of cell-surface biotinylation kits during the COVID-19 pandemic, at least two experimental repeats were conducted on all our stable cell lines – a representative blot is shown in (Figure 5B). However, as the p.Ala1687Pro mutation had not been assessed in the transient expression system, we prioritised running three experimental replicates for this cell line.

**Figure 5:**
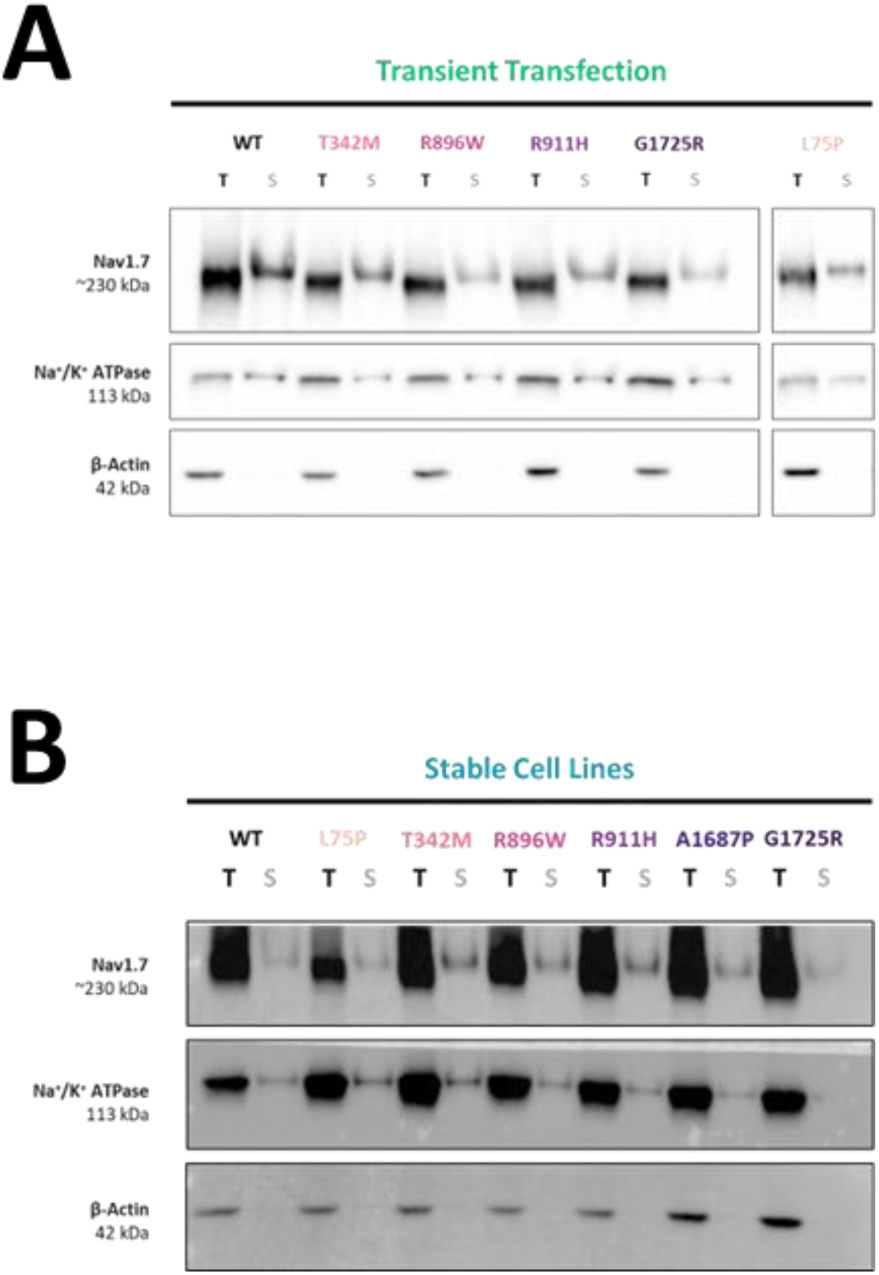
Cell membrane localisation of Na_v_1.7 assessed by cell surface biotinylation. Cell surface biotinylation assays were performed on HEK293 cells expressing Nav1.7. Representative western blots from preliminary experiments using a transient transfection approach (n=3) and in the Na_v_1.7 Jump-In™ T-REx™ HEK293 stable cell lines (n=2) are shown in (A) and (B) respectively. The p.Ala1687Pro mutation was only assessed in the stable cell line system as this patient was diagnosed later in the study. Samples representing total (T) cell lysates and surface (S) localised proteins were probed for Na_v_1.7, as well as Na^+^K^+^ATPase as a positive control and β-actin as a negative control in the surface fractions. These results show that all Na_v_1.7 mutants reached the cell membrane.

The results in our stable cell lines agreed with the earlier findings, indicating none of the mutations caused a defect in forward trafficking to the cell membrane. Taken together with the negligible current density measured in the stable cell lines, our results suggest the six CIP mutations cause localised structural changes sufficient to abolish Na_v_1.7 sodium conductance.

## Discussion

The relationship between Na_v_1.7 mutations and extreme pain phenotypes is well-established (Dib-Hajj and Waxman, 2019). Nevertheless, as genetic testing becomes more accessible, this accelerated data acquisition has led to thousands of variants being detected that require expert interpretation before being causally associated with disease (Schon et al., 1993; Duzkale et al., 2013; MacArthur et al., 2014). In this study, we report on six novel missense *SCN9A* mutations identified from CIP patients. Through a series of molecular assays we showed that, despite intact trafficking to the cell membrane in a HEK293 cell background, all variants led to a failure of sodium conductance. *In silico* analyses suggest the native residues stabilise intramolecular interactions necessary for Na_v_1.7 activation and maintaining the integrity of the ion permeation pathway - excluding the N-terminal residue Leu75 whose functional contribution is not yet understood.

*SCN9A*-CIP is an autosomal recessive condition, with most mutations causing either premature truncation of the protein due to nonsense substitutions, frameshifts or mis-splicing (Goldberg et al., 2007; Shaikh et al., 2018; Marchi et al., 2018), or missense mutations that substitute single functionally critical residues in an otherwise fully folded channel (Cox et al., 2010; Emery et al., 2015; McDermott et al., 2019; Sun et al., 2020). Because patients exhibit such a distinct clinical phenotype, most reported mutations have been associated with disease based on the expected segregation of variants in affected families, population frequency data and bioinformatic predictions (Schon et al., 1993; Duzkale et al., 2013). The new missense *SCN9A* variants detected in our patients met all these requirements; however, the rarity of Na_v_ channel variants does not always correlate with pathogenicity (Waxman et al., 2014). For example, the *SCN9A* variant p.Asn641Tyr was once implicated with epilepsy due to co-segregation with disease in a large American pedigree and allele frequency of 0.001% in the gnomAD database (Singh et al., 2009) but was later found in 18 unaffected individuals of an extended Amish family, contradicting its designation as a pathogenic mutation (Fasham et al., 2020). Reports such as these reinforce the need for laboratory studies to validate the impact of variants on protein function.

Combining the Jump-In™ T-REx™ HEK293 system with our molecular assays allowed us to profile multiple Na_v_1.7 variants in parallel, confirming their pathogenicity. A key benefit of this system was the controlled integration of each *SCN9A* insert (wild-type or mutant) into a single, inducible, pre-determined location of the parental cell genome. Therefore, any significant differences in our assays between the wild-type and mutant channels could be reliably attributed to an effect of the mutation. This contrasts with artefacts caused by differences in the site of genomic integration when using traditional stable cell line approaches, or even differences in transient transfection efficiency. The system could be beneficial for other Na_v_ channelopathies, such as infantile-onset epilepsy, where variant classification and a precise genetic diagnosis can guide early treatment decisions, dramatically improving a patient’s quality of life (Musto et al., 2020; Balestrini et al., 2021).

Electrophysiology remains the gold standard for categorising Na_v_ channel variants, by quantifying any deviations from their normal biophysical profiles that could affect cell excitability and thus human physiology (Glazer et al., 2020; Thompson et al., 2023). These experiments are often conducted in HEK293 cells due to the lack of endogenous Na_v_ channels and ease of use in parallel biochemical assays. In our positive control cell lines, wild-type Na_v_1.7 channels consistently produced peak currents in the 2-3 nA range, while small currents >200 pA could be detected from the previously characterised partial LOF Na_v_1.7 mutant p.Ala1236Glu. In contrast, peak currents in the new CIP cell lines were almost imperceptible, measuring less than the ∼170 pA observed in negative control cell lines lacking Na_v_1.7 protein. These results provide conclusive evidence for LOF, and therefore these missense variants could be classed as pathogenic mutations.

In addition, we used fluorescence microscopy to determine whether the lack of current in the mutant Na_v_1.7 cell lines was caused by defects in trafficking to the cell membrane. While HEK293 cell lines are useful for electrophysiology, the resolution of standard confocal microscopes combined with immunocytochemistry seems insufficient to detect Na_v_1.7 at the cell membrane in this cell background (Sun et al., 2020) – although a “cell membrane ring” has been observed in PC12s and undifferentiated SH-SY5Y cells (Cox et al., 2010; Vetter et al., 2012). In our wild-type and mutant HEK293 cell lines, Na_v_1.7 was expressed diffusely in the cytoplasmic pool to a similar degree, without obvious overlap with an ER marker. Terminally misfolded ion channels tend to form small aggregates retained within the secretory pathway (Manganas et al., 2001; Marangoni et al., 2011), so we inferred our mutations do not affect the overall tertiary structure of Na_v_1.7 channels.

Future studies of Na_v_1.7 trafficking could benefit from recent developments in higher-resolution microscopy methods and fluorescent tagging approaches that have been used to visualise and quantify changes in Na_v_ channel expression at the cell surface of HEK293 cell lines and primary neuronal cultures (Sizova et al., 2020; Tyagi et al., 2024). However, cell-surface biotinylation has been successfully employed to measure changes in Na_v_1.7 expression at the cell membrane (Dustrude et al., 2013; Laedermann et al., 2013) and is less resource-intensive than these techniques. In both transient transfections and in our stable cell lines, we found that all six Na_v_1.7 mutants reached the cell membrane. Combined with our electrophysiological data where no sodium currents were detected, these results suggest that our mutations disrupt confined intramolecular interactions necessary for Na_v_1.7 to undergo conformational changes during voltage-dependent activation and sodium conductance.

Two of our CIP mutations, p.Ala1687Pro and p.Arg911His, are highly likely to have a deleterious impact on the shape of the ‘DEKA’ ion selectivity filter such that the sodium ion permeation pathway is obstructed. The charged D/E/K residues have been proposed to form the sodium ion binding site based on size and electrostatic interactions (Favre et al., 1996), as point substitutions significantly reduce sodium ion selectivity over other metal ions (Heinemann et al., 1992; Schlief et al., 1996; Sun et al., 1997). Alanine has typically been considered the least important residue within the filter due to its small size and lack of charge, although these properties are important alongside the D/E/K residues for the exclusion of divalent calcium ions (Heinemann et al., 1992). The p.Ala1687Pro mutation is relatively conservative in terms of biochemical properties as both residues are small and uncharged. However, proline is known for its inflexibility and for forcing kinks in secondary structures. Furthermore, the homologous mutation in Na_v_1.1 (p.Ala1724Pro) has been reported in a patient with intractable childhood epilepsy; although lacking functional analysis, it is in keeping with the association between *SCN1A* LOF mutations and Dravet syndrome (Wang et al., 2012; Meisler et al., 2021).

The passage of sodium ions through the Na_v_1.7 pore may also be disrupted by the p.Arg911His mutation. This arginine is located four residues above the domain II glutamic acid of the selectivity filter; the proximity and opposite charges of the Arg911 and Glu916 sidechains form a salt bridge that appears to stabilise the orientation of glutamic acid towards the centre of the pore. The p.Arg911His mutation would normally be considered a neutral substitution as both amino acids are equally large and the histidine residue can be positively charged in certain chemical environments (Betts and Russell, 2007). However, it is more likely to be uncharged when exposed to the extracellular fluid with a pH of ∼7.3 (Casey et al., 2010; Requião et al., 2017), which would disrupt the salt bridge with Glu916. Mutations of the homologous arginine in Na_v_1.1 (Arg946) and Na_v_1.2 (Arg937) to histidine or cysteine were also shown to cause a complete loss of conductance (Liao et al., 2010; Volkers et al., 2011; Ben-Shalom et al., 2017), while those in Na_v_1.5 (Arg893) were associated with the cardiac LOF phenotype Brugada syndrome in five unrelated individuals (Kapplinger et al., 2010). These reports underscore the importance of the arginine at this location across the human Na_v_ channel family.

The third CIP mutation, p.Arg896Trp, has been independently characterised by another group by whole-cell patch clamp and our results agreed with their findings of null conductance (McDermott et al., 2019). Another CIP patient has also been reported with a different missense substitution at the same location, p.Arg896Gln, shown to be non-conducting in HEK293 cells and displayed a reduced capacity to reach the plasma membrane in PC12 cells when assessed by fixed-cell confocal microscopy (Cox et al., 2010). Mutations of the homologous arginine in Na_v_1.5 (Arg878) to cysteine or histidine have been reported in Brugada syndrome and sick sinus syndrome patients. Functional analysis established the p.Arg878Cys mutant is non-conducting and excluded trafficking defects (Zhang et al., 2008; Gui et al., 2010; Kapplinger et al., 2010). These mutations point again to a critical role of the native arginine residue. Given its sidechain bridges the pore loops of domains II and III and forms a salt bridge with Asp1406, another fully conserved residue in across all Na_v_1.7 homologues, this arginine may be involved in stabilising the extracellular vestibule to optimally guide solvated sodium ions towards the pore.

The structural impacts of the next two CIP mutations on Na_v_1.7 function, p.Thr342Met and p.Gly1725Arg, are less clear. Thr342 is located on the DI:S5-S6 pore loop, although it is further from the permeation pathway than the previously discussed mutations. In Na_v_1.5, p.Thr351Ile was found to cause misfolding which led to a reduction of membrane-localised channels and ∼75% reduction in peak whole-cell Na^+^ currents in HEK293 cells; the trafficking defect was partially corrected *in vitro* by incubation with mexiletine and resulted in measurable whole-cell currents (Pfahnl et al., 2007). In Na_v_1.1, p.Thr363Pro and p.Thr363Arg were also associated with Dravet syndrome phenotypes, although the mechanism and magnitude of LOF were not investigated (Zuberi et al., 2011; Wang et al., 2012). While the loss of this threonine across human Na_v_ channels may be universally pathogenic, the complete loss of conductance and absence of a trafficking defect caused by the Na_v_1.7 p.Thr342Met mutation suggests that the nature of LOF may depend on the properties of the substituted amino acid.

A similar pattern exists for Na_v_ channelopathies caused by substitutions of the glycine homologous to Gly1725 in Na_v_1.7, located on the top of the DIV:S6 transmembrane segment. While the Na_v_1.7 CIP mutation p.Gly1725Arg is non-conducting (this study and (McDermott et al., 2019)), the Na_v_1.5 Gly1748Asp mutant associated with Brugada syndrome exhibited an >80% reduction in peak current due to a trafficking defect, as well as substantial changes to multiple voltage-dependent parameters (Núñez et al., 2013). An alternative substitution in Na_v_1.1 (p.Gly1762Glu) has been reported in a patient with a sub-phenotype of Dravet syndrome but lacks functional data (Mancardi et al., 2006). The native glycine residue is small and uncharged, conferring high conformational flexibility to this part of the protein. Therefore, substitutions that change glycine to any of these larger, charged amino acids are understandably disruptive. In the case of Na_v_1.7, the p.Gly1725Arg mutation may lock the DIV:S6 segment in place, preventing the radial turn necessary to open the intracellular activation gate (Yang et al., 2013).

Our final *SCN9A*-CIP mutation, p.Leu75Pro, is in the intracellular N-terminal tail. It is the first N-terminal mutation to be reported, across the human Na_v_ channel family, that completely abolishes channel function. It is unclear how this region contributes to Na_v_ channel gating and conductance, however, the association of >40 N-terminal missense mutations in Na_v_1.1 and Na_v_1.5 with a range of LOF and GOF disorders, supports its functional importance (Huang et al., 2017). Of the few N-terminal missense mutations that have been biophysically characterised, p.Arg99His in Na_v_1.7 significantly reduced current density without changes to voltage-dependent properties (Sun et al., 2020), whereas p.Asp12Asn and p.Asp82Gly in Na_v_1.2 reduced channel availability by altering the kinetics of inactivation (Ben-Shalom et al., 2017). In contrast, a Na_v_1.7 GOF mutation, p.Q10R, hyperpolarised voltage-dependence of activation resulting in inherited erythromelalgia (Han et al., 2009).

The N-terminal region was recently resolved in crystal structures of human Na_v_1.7 (Huang et al., 2022). The N-terminus appears to possess some secondary structural elements forming a unit that docks below domain I. Our mutation is found in a highly conserved motif “PLEDLDPYY” which forms a small loop with the side chains of positively charged (glutamic acid, aspartic acid) and hydrophobic (leucine, phenylalanine, tyrosine) amino acids pointed in alternating directions. The p.Leu75Pro substitution could force a premature tight turn in this loop, adjusting the orientation of these residues. The exact function of this motif and the N-terminal tail unit remains to be explored, but hypotheses include a role in channel activation or interactions with a protein binding partner necessary for membrane tethering.

Na_v_ channels are critical to electrical signalling in many physiological systems, and prime targets for therapeutic modulation in disorders of excitability. From Hodgkin and Huxley’s theoretical model of the m- and h-gates controlling a voltage-sensitive and sodium-selective channel (Hodgkin and Huxley, 1952); to Hille and Armstrong delineating the biochemical properties of the transmembrane pore (Armstrong and Hille, 1998); to decades of structural studies illuminating the nuances of how individual residues, segments and domains move independently and in concert to cycle through distinct functional states (Catterall et al., 2020) – we are slowly uncovering the intricacies of these molecular machines. Our study demonstrates both the clinical and fundamental utility of assessing human disease variants, to improve diagnosis and patient care, as well as the potential to yield profound insights into the complexity of protein function.

## List of Abbreviations

CIP: Congenital Insensitivity to Pain
DEKA: D-E-K-A amino acids forming the voltage-gated sodium ion selectivity filter
DI to DIV: Domains 1 to 4 of Na_v_ channels
ER: Endoplasmic Reticulum
GOF: Gain-of-Function
LOF: Loss-of-Function
S1 to S6: Segments 1 to 6 of each Na_v_ channel domain
11L: Exon 11 Long splice variant of Na_v_1.7

## Data Availability Statement

The data generated from this project are available from corresponding authors upon reasonable request.

## Acknowledgements

This project was made possible through a BBSRC iCASE PhD studentship (Project Reference 1801406) and continued support from the UKRI Advanced Pain Discovery Platform and NIHR Cambridge Biomedical Campus award 2021. We also thank Matthew Gratian and Mark Bowen of the Microscopy Facility at Cambridge Institute for Medical Research for their assistance in confocal imaging, and Randy Read, Alexandre Faille and Vasileios Kargas for their advice on structural visualisations.

## Author Contributions

C Geoffrey Woods and John Linley conceptualised and supervised this study. Fiona Cusdin provided key resources. Nivedita Sarveswaran, Ichrak Drissi, Samiha Shaikh and Mike Nahorski developed methodology, performed experiments and reviewed analyses. Nivedita Sarveswaran prepared the manuscript and visualisations. All authors read and approved the final manuscript.

